# Neural representation strength of predicted category features biases decision behavior

**DOI:** 10.1101/2023.05.05.539587

**Authors:** Yuening Yan, Jiayu Zhan, Oliver Garrod, Xuan Cui, Robin A.A. Ince, Philippe G. Schyns

## Abstract

Theories of prediction-for-perception propose that the brain predicts the information contents of upcoming stimuli to facilitate their perceptual categorization. A mechanistic understanding should therefore address where, when, and how the brain predicts the stimulus features that change behavior. However, typical approaches do not address these predicted stimulus features. Instead, multivariate classifiers are trained to contrast the bottom-up patterns of neural activity between two stimulus categories. These classifiers then quantify top-down predictions as reactivations of the category contrast. However, a category-contrast cannot quantify the features reactivated for each category–which might be from either category, or both. To study the predicted category-features, we randomly sampled features of stimuli that afford two categorical perceptions and trained multivariate classifiers to discriminate the features specific to each. In a cueing design, we show where, when and how trial-by-trial category-feature reactivation strength directly biases decision behavior, transforming our conceptual and mechanistic understanding of prediction-for-perception.

## Introduction

In vision science, Helmholtz’s “unconscious inferences” emphasized that perception not only relies on the bottom-up visual input, but also on top-down expectations of what this input might be^1–3^. Visual expectations should predict upcoming visual contents, thereby facilitating their processing from the input^4–6^ for perceptual decision behavior^7,8^. However, two related challenges must be addressed to develop better specified theories and models of prediction-for-perception: (1) What are the specific contents that are predicted about a visual category and (2) how do these predicted contents influence behavior?

Predictions of visual information about a category arise from memory^9^. For example, to predict faces and cars, current theories and models propose that the visual features that represent faces (e.g., eyes and nose) and cars (e.g., wheels and windshield) in memory propagate down a hierarchy of brain regions^10,11^, from the hippocampus to the lower levels of sensory cortex^12,13^, under pre-frontal control^14–16^. To access these predictions in the visual hierarchy, researchers typically use a two-stage analysis. First, they train multivariate binary *category-contrast* classifiers to discriminate the bottom-up neural response (i.e. M/EEG or fMRI responses) to pairs of stimulus categories. Then, they reuse these bottom-up classifiers to decode the top-down cued-reactivation of the pattern of neural activity that represents the predicted category-contrast before the stimulus is shown^10,11,17,18^.

Much has been gained from this approach to understand predictions, from low-level perceptual contrasts (e.g. photographs vs. drawings) to semantic contrasts (animate vs. inanimate)^10,19^. Importantly, contrast reactivations from memory can reveal top-down, semantic-to-perceptual effects^10^, similar to those reported in mental imagery^11^. That is, the prediction process reverses in time the bottom-up representations of stimuli in the occipito-ventral pathway by converting mnemonic categories (in hippocampus) to perceptual representations (in occipital cortex)^20^. Furthermore, predictions of perceptual contrast can sharpen the representations of subsequent stimuli, from occipital cortex (e.g. for Low vs. High Spatial Frequencies, SF)^5^ to medial prefrontal cortex (e.g. for higher-level “face vs. scene vs. object”)^6^.

Notwithstanding these important findings, it is important to realize that studying predictions of such category contrasts cannot address the specific visual information contents that the brain predicts about each category, and how these contents influence subsequent decisions. For example, reactivation of a contrast that discriminates the categories of faces and cars could be driven either by car features (e.g. wheels vs. no wheels), or by face features (eyes vs. no eyes), or by features from both categories (e.g. wheels vs. nose). Typical category-contrast classifiers cannot distinguish these cases because they cannot quantify the predictions of the visual features specific to each category (presence of wheels or a nose); they can only quantify that some (typically unspecified) features differ between the categories. This shortcoming reduces the resolution at which the process of predicting specific perceptual information from memory can be quantitatively described and therefore hinders studies of its mechanistic understanding.

Here, to address the challenge of specifying visual predictions, we trained linear *category-feature classifiers* to decode the representational strength of the specific visual features that define each category. To preview our key results, we show that individual participant, trial-by-trial reactivations of these predicted category-features do bias psychometrically measured decision behavior–and do so more reliably and more strongly than the reactivations of the typical category-contrast.

## Results

We used a realistic stimulus known to drive mutually exclusive perceptions based on two visual contents represented within different Spatial Frequency (SF) bands^21^. Dali’s *Slave Market with the Disappearing Bust of Voltaire* can be perceived either as the bust of “Voltaire” or as two “Nuns” (Figure 1A, Original) from respectively processing Low (LSF, 8 cycles per image) or High image SF (HSF, 16 and 32 cycles per image)^21^, Figure 1A, Features–see *Methods, Stimuli*.

**Figure 1.**
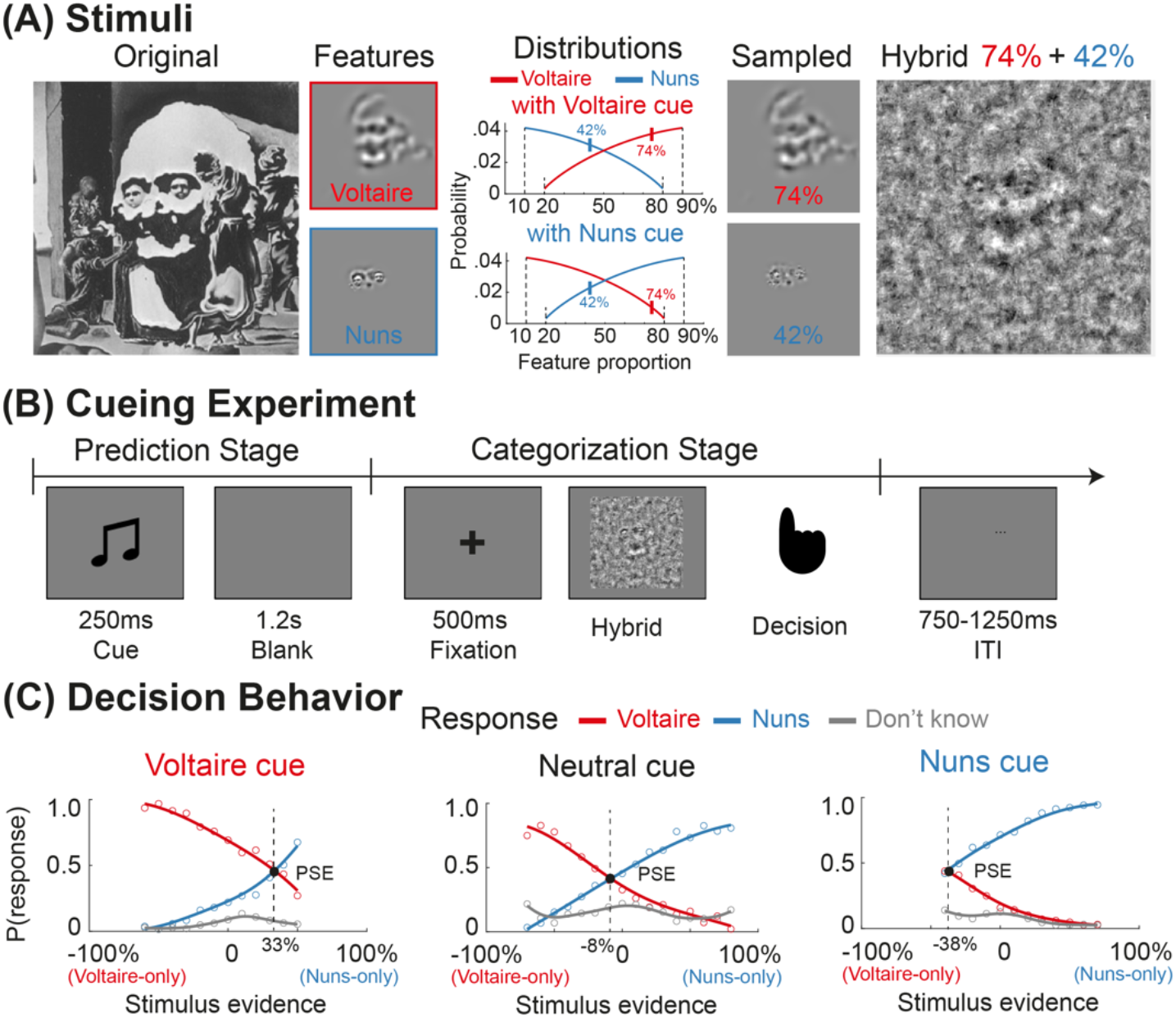
Experimental design and behavioral result. **(A) Stimuli**. From the <u>Original</u> ambiguous stimulus, we applied filters to extract the Voltaire and Nuns stimulus <u>Features</u>^21^, respectively represented within LSF 8 cycles/image and HSF 16 to 32 cycles/image, see *Methods, Stimuli*. On each trial, a hybrid stimulus comprised randomly and independently selected percentages of the Voltaire (red) + Nuns (blue) features filtered with spatial Gabors. These proportions were independently and randomly sampled from the <u>Distributions,</u> based on whether the cue was Voltaire vs. Nuns (under neutral cueing, proportions were random between 0% and 100%). The <u>Hybrid</u> example illustrates proportions of 74% Voltaire + 42% Nuns Gabor features (see vertical marks in the distribution) inserted in Gabor background noise, see *Methods, Stimuli*. **(B) Cueing Experiment**. *<u>Prediction</u> <u>Stage</u>*: A 250ms pure tone cued either the Voltaire (196Hz) or Nuns (1760Hz) stimulus distribution, or was neutral (880Hz), followed by a 1.2s blank interval. *<u>Categorization</u> <u>Stage</u>*: A 500ms fixation was followed by a hybrid image that remained on the screen until response (“Nuns” vs. “Voltaire” vs. “Don’t Know”, 3-AFC), followed by a 750 to 1250ms inter-trial interval (ITI) with jitter. **(C) Decision Behavior**. Relationships between stimulus feature evidence and response probability–i.e. color-coded curves of P(“Voltaire”), P(“Nuns”) and P(“Don’t Know”), for each auditory cue (panel). The Point of Subjective Equality (PSE, black dot) is the level of stimulus evidence of equally likely P(“Voltaire”) and P(“Nuns”). <u>Left and right panels</u> show that Voltaire and Nuns cues shift the PSE in opposite directions compared with the neutral cue (<u>central panel</u>).

Prior to the experiment, we trained participants (*N*=10) to couple auditory cues (a 250ms pure tone at 196Hz vs. 1760Hz) with the Voltaire vs. Nuns features that cause each perception. We also included a neutral 880Hz auditory cue that had no predictive value. In the experiment, participants saw hybrid images that comprised varying proportions of LSF “Voltaire” vs. HSF “Nuns” Gabor features added to a noise background. Critically, these proportions of Voltaire and Nuns Gabor features were independently and randomly sampled on each trial from the pre-defined distributions shown in Figure 1A–illustrated with a hybrid stimulus that adds a random selection of 74% of the Gabors underlying perception of Voltaire to a random selection of 42% of the Gabors underlying perception of the Nuns.

Each experimental trial comprised a Prediction Stage followed by a Categorization Stage (Figure 1B). An auditory cue predicted the features of the upcoming stimulus (Figure 1A). We instructed participants to report their strongest perception: “Voltaire,” vs. “Nuns,” vs. “don’t know”. Our analyses then traced where, when, and how the auditory cues reactivate from the participant’s memory the visual representations of specific Voltaire and Nuns Gabor features that influence behavior (3,375 trials/participant, decoding and statistical inference performed within each participant^22,23^). We compared these novel reactivations of category-features (of Voltaire and Nuns) to the classic reactivations of category-contrast (of Voltaire vs. Nuns).

### Cued-Predictions Bias Perceptual Decisions

Figure 1C confirms that predictive cues changed perceptual decision behavior. To understand this change, we plotted group-level psychometric curves. Separately for each predictive auditory cue (panel) and each possible decision response (colored curves), we computed the relationships between stimulus feature evidence (relative percentages of sampled Voltaire vs. Nuns stimulus Gabor features, X axis) and response probability (Y axis)–see *Methods, Cued-Predictions bias perceptual decisions*. The curves for Voltaire and Nuns responses intersect at the Point of Subjective Equality (PSE), which indicates the level of stimulus evidence (i.e. X axis of relative proportions of Voltaire and Nuns Gabor feature evidence) for which Voltaire and Nuns responses (Y axis, probabilities) are equally likely. Under the neutral cue (Figure 1C, central panel), the PSE is -8%. That is, 8% more Voltaire than Nuns evidence is required for equiprobable Voltaire and Nuns responses, a small bias towards Nuns. Cueing Voltaire (left panel) shifts the PSE to 33% more Nuns evidence. Cueing the Nuns (middle panel) shifts the PSE to 38% more Voltaire evidence. We replicated these cue-induced shifts of the PSE in each participant, leading to the conclusion that predictive cueing strongly biases responses towards the cued behavior. Supplemental Table S1 reports individual participants PSE shifts across cues; Supplemental Figure S1 shows the bias effect across the full range of feature evidence in individual participants.

### Dynamic predictions of category-contrast vs. category-features in the brain

To understand where, when, and how the auditory cues of Voltaire and Nuns reactivate the predicted visual contents, we applied the analyses summarized in Figure 2. First, we trained the typical category-contrast classifiers (Figure 2A) to discriminate 100% Voltaire from 100% Nuns images, using each participant’s bottom-up MEG response data from a localizer run prior to the cueing experiment (passive viewing). At the Prediction Stage (Figure 1B), we applied these classifiers to quantify the cued reactivations (i.e. prediction strength) of the Voltaire vs. Nuns category-contrast. We then tested how per-trial prediction strength influences behavior (Figure 2C). As discussed, category-contrast classifiers cannot quantify separately the reactivation of the features specific to each category. We therefore trained separate category-feature classifiers to learn the relationship between Voltaire- and Nuns-specific Gabor features and brain responses (at the Categorization Stage, under neutral cueing, Figure 2B). We applied these classifiers at the Prediction Stage to quantify the cued-reactivations of the category-specific features–i.e., their predictions (Figure 2C). Then we compared how per-trial category-contrast and category-feature prediction strength influences behavior. To preview our results, category-feature predictions are better localized, more specifically driven by the auditory cues, with per-trial reactivation strength that biases participant’s behavior more.

**Figure 2.**
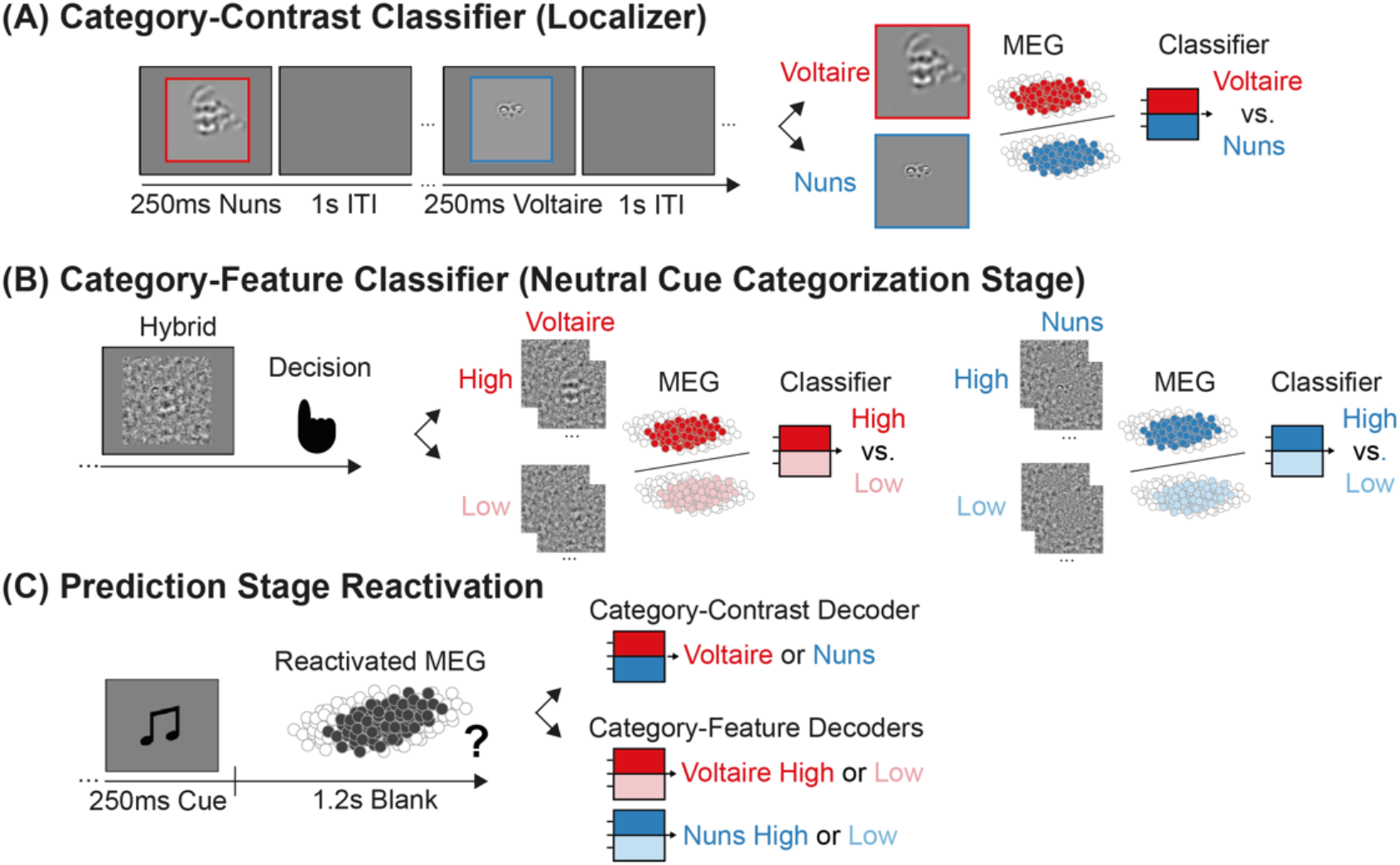
Prediction Reactivation. **(A) Category-Contrast Classifier**. Trained on a localizer run prior to the cueing experiment, category-contrast classifiers learn to discriminate the bottom-up patterns of sensor-level neural responses to each stimulus category (color-coded as Voltaire vs. Nuns and measured with MEG). **(B) Category-Feature Classifier**. Trained on Categorization Stage sensor-level data under the neutral cue, category-feature classifiers learn to discriminate the High (>70%) vs. Low (< 30%) proportions of Voltaire Gabor features and separately Nuns Gabor features. **(C) Prediction Stage Reactivation**. Following the auditory cues that predict the Voltaire vs. Nuns Gabor feature distributions (Figure 1A), the category-contrast and the category-feature classifiers quantify the single-trial reactivation strength of the sensor-level pattern of MEG activity every 4ms of the Prediction Stage.

### Prediction reactivation with category-contrast classifiers

Previous studies suggested that top-down predictions of the category-contrast reverse their bottom-up flow^10,11^. We therefore applied category-contrast classifiers to the Prediction Stage of our experiment to establish a baseline of results to compare against our new category-feature classifiers.

In each participant, we trained and cross-validated binary category-contrast classifiers on images of 100% Voltaire and 100% Nuns, every 4ms of the 0 to 400ms post-stimulus response of the MEG localizer (Figure 2A). We similarly trained classifiers in a separate MEG localizer of the auditory cues (196 Hz, Voltaire vs. 1760 Hz, Nuns). We ran both localizers before the main cueing experiment to avoid contamination (see Supplemental Figure S2 and *Methods, Visual and Auditory localizers*). We further split the classifiers into the Early set, trained from 75 to 150ms post stimulus (to cover the early P1 Event-Related Potential, ERP, time window associated with processing low-level visual features^24,25^), and the Late set, trained from 150 to 280ms (to cover the later N170 and N250 ERPs associated with higher-level categorization^26–32^), see *Methods, Localizer cross-validation*.

We applied these Early and Late bottom-up category-contrast classifiers at the top-down Prediction Stage of the cueing experiment (illustrated in Figure 2C; Figure 1B). This analysis produced, every 4ms, per-trial classifier decision values that quantify the prediction strength of the category-contrast. To quantify the time course of prediction strength across trials, for each classifier we computed MI(ground truth Voltaire vs. Nuns cue; decision value_*t*_,)–FWER corrected over 100 localizer training time points and 150 Prediction Stage testing time points, *p*<0.05, one-tailed, see *Methods, Category-contrast reactivation*. We then selected two classifiers: That with maximum prediction reactivation from the Early set (75 to 150ms); that with maximum prediction reactivation from the late set (150 to 280ms)–i.e. both selected from the matrix of localizer training time points × Prediction testing time points. The performance of these two classifiers quantifies how strongly the auditory cues reactivate the dynamic prediction of the Voltaire vs. Nuns category-contrast.

In Figure 3, dark purple curves show across participants that Late classifiers patterns are more strongly reactivated by the cue than than Early classifiers (light purple). We localized the MEG sources that represent this prediction–i.e. by computing MI(Late classifier decision value; MEG source activity) on all 8,196 individual sources at Prediction. We found initial ventral stream involvement from temporal cortex (MTG, STG, PHC, FG) down to occipital cortex (LG, PCAL, CUN, LOC), followed by bi-lateral occipital cortex. Thus, top-down predictions of the Voltaire vs. Nuns category-contrast reverse the bottom-up flow of its visual processing ^10,11^. We replicated the visual category-contrast predictions in 9/10 participants–FWER, *p*<0.05, one-tailed, Bayesian population prevalence (BPP) ^22,23^, with maximum a posteriori probability (MAP) estimate of the population prevalence of the effect of 9/10 replications = 0.9 (95% highest posterior density interval, HPDI [0.61 0.99]), see Supplemental Figure S3 for individual participant results.

**Figure 3.**
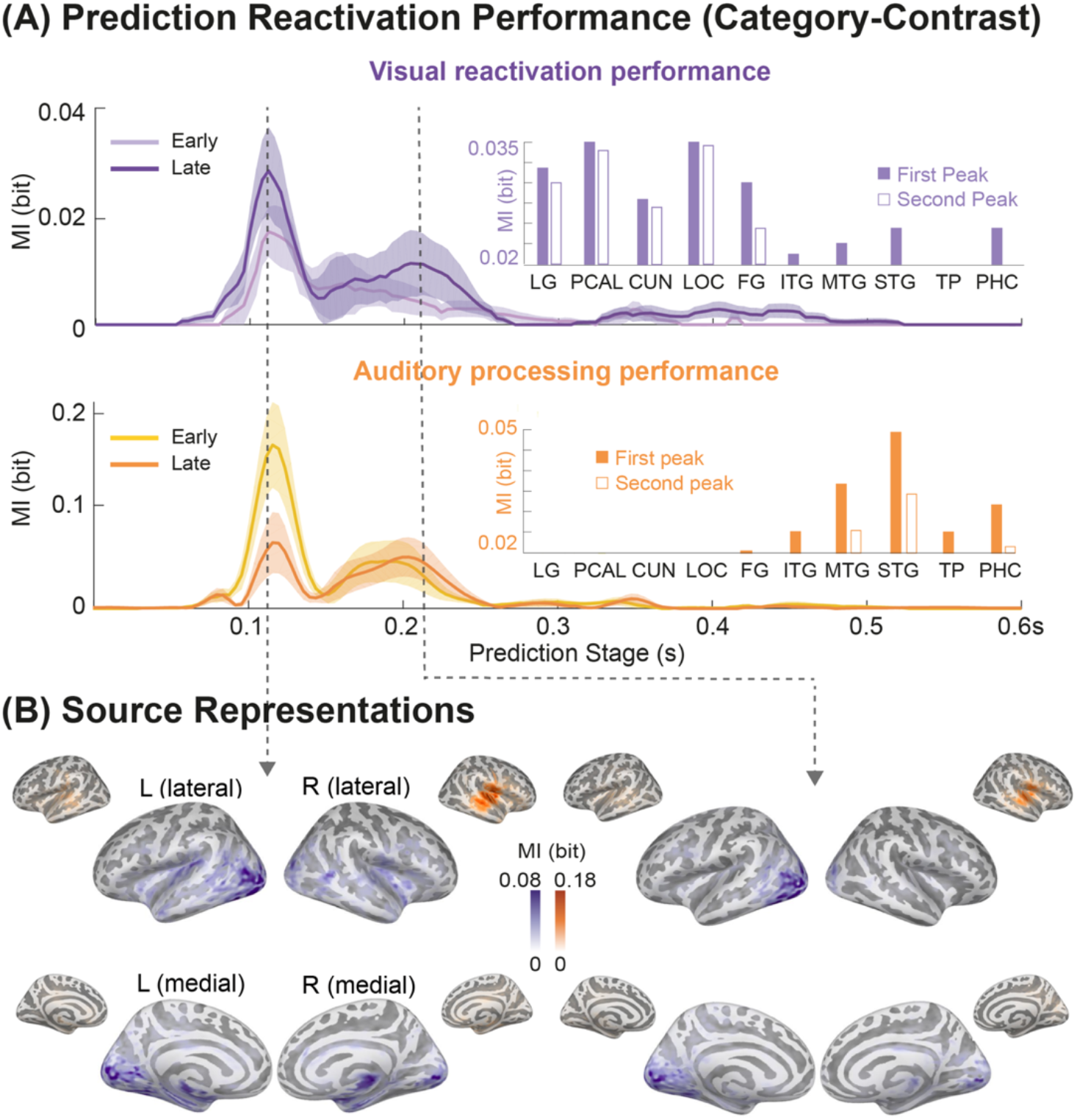
Voltaire vs. Nuns Category-Contrast Reactivation. **(A) Prediction Reactivation Performance**. <u>Purple curves</u> show the dynamic prediction reactivation performance of the Voltaire vs. Nuns category-contrast at each time point of the Prediction Stage, for the Early classifier (with the highest performance over 75-150ms post-cue, light purple), and Late classifier (with highest performance over 150-280ms post-cue, dark purple) averaged across participants–shaded regions denote ± standard errors of the mean. To control for the propagation of the auditory cue contrast, we trained auditory classifiers to discriminate the two cues on the auditory localizer data and similarly tested their decoding performance on the same Prediction Stage responses (orange). The curves show that the late bottom-up classifier (dark purple) better models the two peaks of Voltaire vs. Nuns visual prediction reactivation (dashed arrows) whereas the early classifier (light orange) better models the two peaks of auditory decoding. **(B) Source Representations**. <u>Cortical surface maps</u> reveal the MEG sources that contribute to the two reactivation peaks of the Late category-contrast classifier (dark purple) and Early auditory classifier (light orange), computed as MI(classifier decision value; MEG source activity). <u>Bar plots</u> in **(A)** show their mean representation strength (MI) across all sources in each ROI at the two peak time points (Early: filled bars; Late: unfilled bars). Abbreviations: Lingual gyrus (LG), pericalcarine cortex (PCAL), cuneus (CUN), lateral occipital cortex (LOC), fusiform gyrus (FG), inferior temporal gyrus (ITG), middle temporal gyrus (MTG), superior temporal gyrus (STG), temporal lobe (TP), parahippocampal cortex (PHC).

We controlled the distinct propagation of the auditory cue contrast by decoding the Prediction Stage with classifiers trained on the auditory localizer data (orange). Initially discriminated in temporal lobe (ITG, MTG, STG, TP and PHC 75-150ms post-cue), the auditory cue contrast does not propagate beyond MTG and STG. Thus, the auditory cue contrast and the reactivated visual category contrast propagate differently–see brain regions histograms and *Methods, Source representation of category-contrast*. We replicated this result in all participants (BPP = 1.0 [0.75 1.0]). See Supplemental Figure S3 for individual participant results.

### Prediction reactivation with category-feature classifiers

We now turn to the reactivations of the specific Gabor features of each predicted category. Separately for Voltaire and Nuns, we grouped stimuli presented under the neural cue into those with High (> 70%) vs. Low (< 30%) Gabor features contents (see Figure 1A). We trained new bottom-up category-feature classifiers (Figure 2B) to discriminate High vs. Low Gabor features from MEG sensor responses (every 4ms of the Categorization Stage, under the neutral cue, see Methods, *Category-feature classifier cross-validation*). We used these category-feature classifiers to quantify the reactivations of the Gabor features of Voltaire and Nuns at Prediction, under Voltaire and Nuns cueing–i.e. separately computing MI(Voltaire vs. neutral cue; Voltaire-feature-classifier decision value) and MI(Nuns vs. neutral cue; Nuns-feature-classifier decision value). We further selected the best performing classifiers at the peak time point of Prediction decoding–FWER corrected over 100 Categorization Stage training time points and 150 Prediction Stage testing time points, *p*<0.05, one-tailed, see *Methods, Category-feature reactivation*. Here, we replicated the significant Prediction Stage decoding (FWER corrected over 100 Categorization Stage training time points and 150 Prediction Stage testing time points, *p*<0.05, one-tailed) in 8/10 participants for Voltaire (BPP = 0.8 [0.49 0.96] (MAP [95% HPDI]) and 9/10 participants (BPP = 0.9 [0.61 0.99] (MAP [95% HPDI]) for Nuns, see Supplemental Figure S4 for individual results).

We then localized the brain sources of these category-feature predictions, computing MI(Voltaire-feature-classifier decision value; MEG source activity) and MI(Nuns-feature-classifier decision value; MEG source activity), on 8,196 sources, at peak reactivation time, see *Methods, Category-feature reactivation*. Figure 4A shows that category-feature predictions comprise bilateral PHC, but also that they lateralize to left fusiform gyrus (FG), lingual gyrus (LG), pericalcarine cortex (PCAL) and lateral occipital cortex (LOC) for coarser-scale Voltaire, but bilateral early visual cortex and right LOC for finer-scale Nuns^33,34^. Thus, category-feature classifiers lateralized the reactivations of the predicted features, a result congruent with lateralized bottom-up representations of scale information^33,34^.

**Figure 4.**
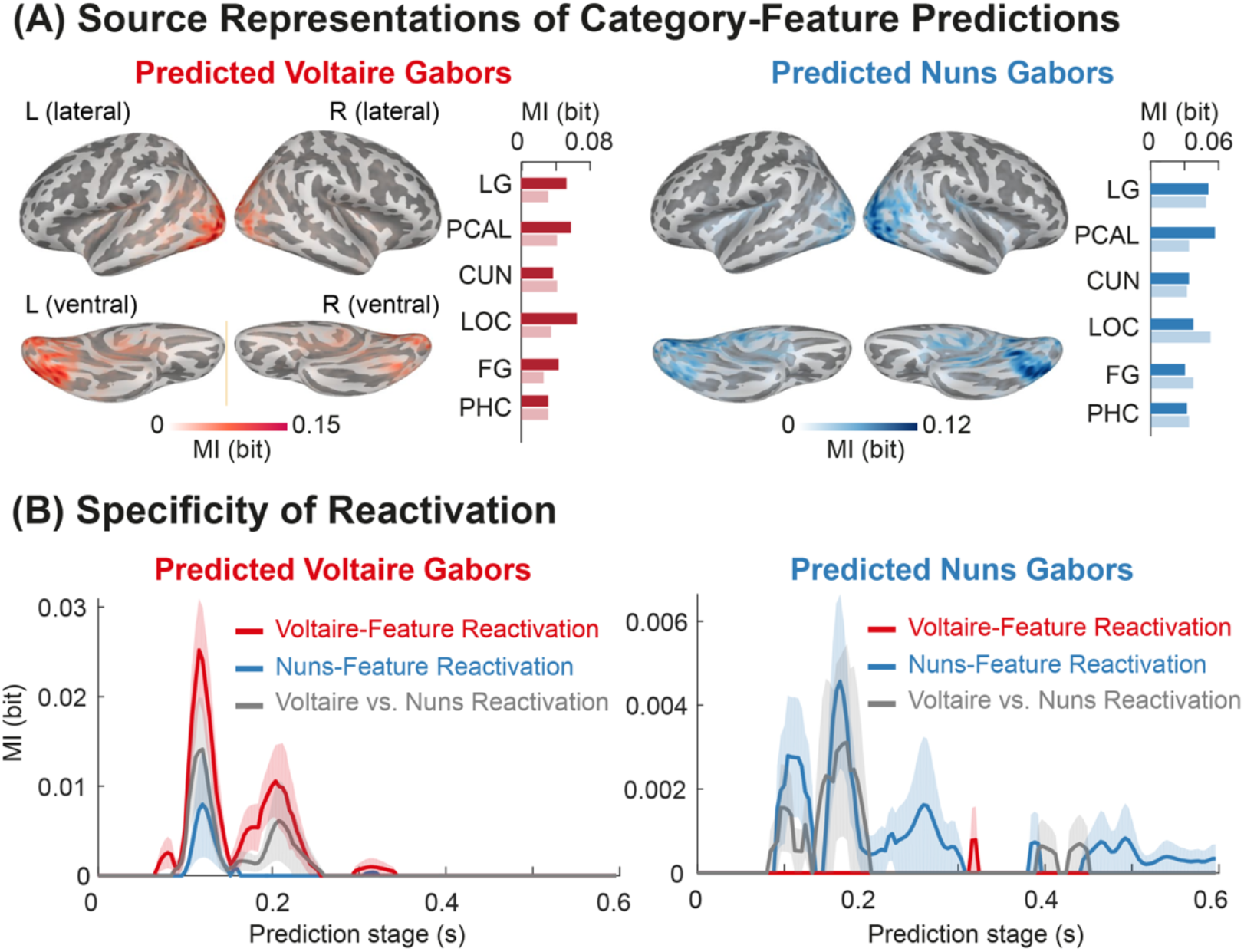
Predictions of Voltaire and Nuns Gabor Features. **(A) Source Representations**. <u>Cortical surface plots</u> localize the MEG sources that contribute to category-feature predictions, at the maximal time point of cued-reactivation of the pattern that represents the discrimination of High vs. Low Gabor features for Voltaire (red) and for Nuns (blue). <u>Bar plots</u> indicate mean (across participants) reactivated category-feature prediction strength (MI) at peak time, in left (darker) and right (lighter) hemispheres, showing that (1) bilateral PHC represent Voltaire and Nuns Gabor predictions; (2) Voltaire Gabor predictions are left lateralized (in FG, LG, PCAL, LOC); Nuns Gabor predictions are right lateralized (in LOC). **(B) Specificity of reactivation**. <u>Color-coded curves</u> show the relative performance of the best category-feature (red for High vs. Low Voltaire Gabors; blue for Nuns) and best category-contrast (Voltaire vs. Nuns) classifiers during the Prediction Stage.

### Specificity of cued reactivations

To investigate how selectively auditory cues reactivate visual predictions, we contrasted reactivations when cues are Voltaire vs. neutral and when they are Nuns vs. neutral. For both contrasts, we compared the performance of the category-contrast and category-feature classifiers (see Figure 4B and *Methods, Specificity of cued-reactivations*). For each feature, we found selective reactivation for the cue contrast is highest with the category-feature classifier, indicating a higher cue-specificity. That is, the Voltaire cue contrast reactivates the Voltaire-feature classifier more strongly than the Nuns-feature classifier and category-contrast classifier. We replicated this cue-specificity in 8/10 participants of for Voltaire cued reactivations (Bayesian population prevalence = 0.8 [0.49 0.96]; MAP [95% HPDI]) and 7/10 participants for of Nuns cued reactivations (Bayesian population prevalence = 0.7 [0.53 0.82]; MAP [95% HPDI]). Thus, the greater specificity of cued-reactivations of the category-feature classifiers provides a finer conceptual resolution to study the mechanisms that predict the visual contents of a category that the category-contrast classifiers.

### Reactivation of category-features at Prediction bias behavior at Categorization

Finally, we compare how the per-trial, cued-reactivations (of category-contrast vs. category-feature) in the brain during the Prediction Stage change the response probabilities of decision behavior at the subsequent Categorization Stage. Specifically, we examined how the top 30% (strong) vs. bottom 30% (weak) trials of classifier reactivations change the psychometric relationship between stimulus evidence and response probabilities (see *Methods, Reactivation biases behavior*). We report how category-feature and category-contrast reactivations change response in Figure 5A and 5B, respectively.

**Figure 5.**
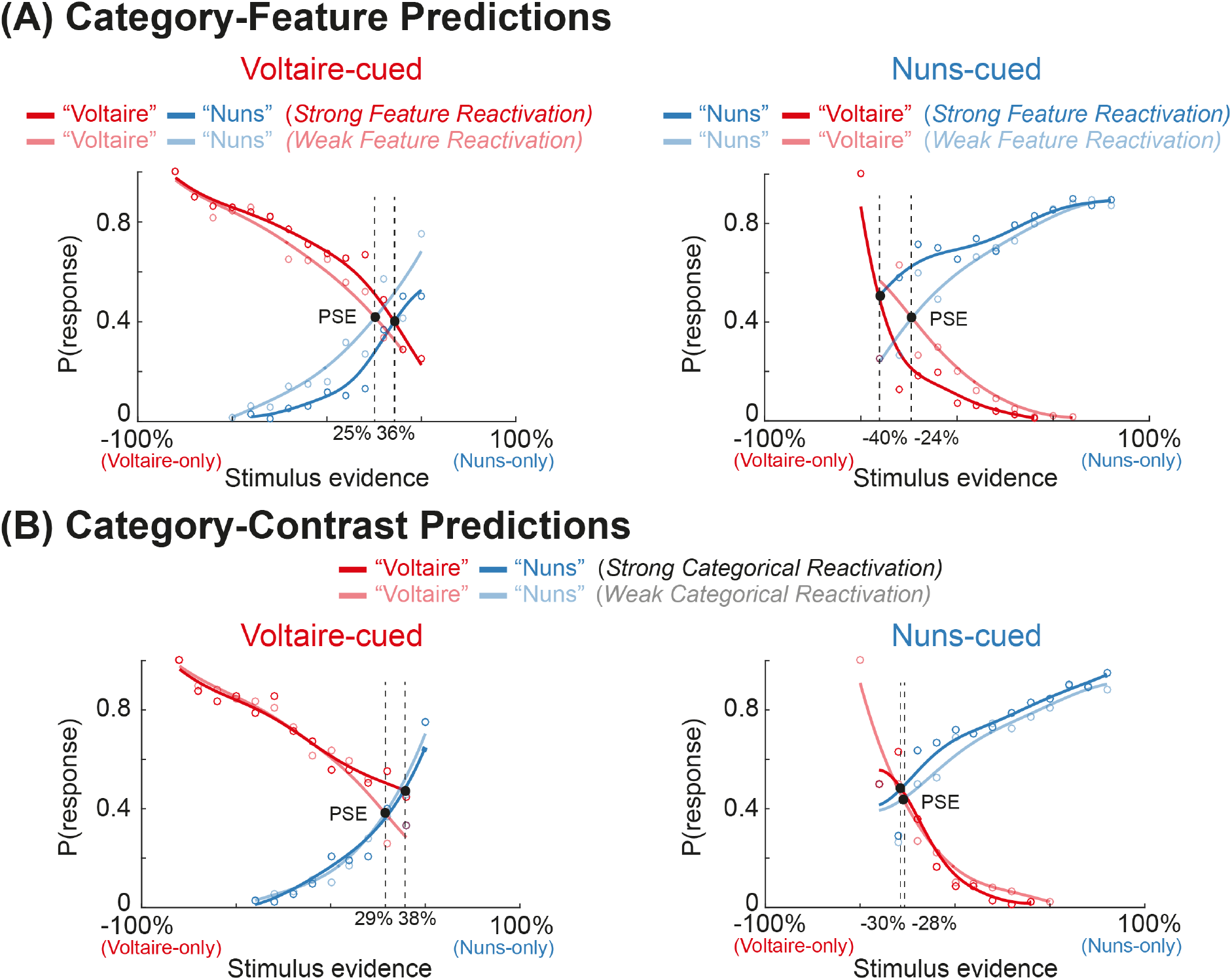
**(A) Category-Feature Predictions**. For Voltaire- and Nuns-cued trials (panels), we independently computed the relationships between stimulus feature evidence (i.e. percentages of Voltaire vs. Nuns Gabor features, X axis), response probabilities (i.e., P(“Voltaire”) (red curves), P(“Nuns”) (blue curves), Y axis) for different strengths of predicted category–i.e., Voltaire-cued (left panel), Nuns-cued (right panel) and strong (opaque curves) vs. weak (transparent curves) prediction reactivation strengths of the category-feature decoders. The black dot is the PSE of equal P(“Voltaire”) and P(“Nuns”). The curves reveal response biases with response bias (vertical offset of light vs dark curves) across a wide range of Gabor feature evidence. The differences in the PSE between the strong and weak reactivation conditions quantifies this bias. **(B) Category-Contrast Predictions**. Similar analyses with the category-contrast decoder of Voltaire vs. Nuns only weakly biases perceptual decisions, and only at high levels of Gabor feature evidence.

Figure 5A shows that strong vs. weak category-feature reactivations at prediction bias decision behavior. Specifically, they shift the PSE by 11% (25% to 36%) on Voltaire-cued trials and -16% (−24% to -40%) on Nuns-cued trials. We replicated these significant PSE shifts (FWER corrected over corrected over 100 Categorization Stage training time points and 150 Prediction Stage testing time points, *p*<0.05, two-tailed) in 8/10 participants (Bayesian population prevalence = 0.8 [0.49 0.96] (MAP [95% HPDI]), see Supplemental Table S2 for the individual results. In comparison, Figure 5B shows that category-contrast reactivations (Nuns vs. Voltaire, trained on 100% localizer stimuli), shift the PSE by only -2% (Nuns predictions, only 2/10 participants with significant effect) and 9% (Voltaire, only 3/10 participants with significant effect), see Supplemental Table S3 for the individual results.

In sum, the per-trial prediction strength of category-features biases the participant’s decisions under both cues. Notably, the magnitude of the bias is comparable in size to the full effect on behavior when the cue is fixed. That is, even with a fixed Voltaire or Nuns cue, the changes in PSE due to reactivations of category-specific features are 27% (Voltaire) and 52% (Nuns) of the changes in PSE that are due to the cues themselves–i.e., category-feature reactivation under the Voltaire cue changes PSE by 11%, whereas the behavioral effect of the Voltaire cue vs. the neutral cue was a PSE change of 41%; For the Nuns cue numbers are -16% (reactivation under Nuns cue) vs. -30% (Nuns vs neutral cue). The effect of category-feature reactivations on behavior is stronger and across a wider range of stimulus evidence compared to the category-contrast classifier. Thus, focusing on category-specific feature predictions can disentangle the decoded category-contrast to improve our understanding of the mechanisms of prediction, from cue-reactivations of category-specific features to the effect of these predictions on behavior.

## Discussion

To develop prediction-for-perception, we must access the dynamic perceptual contents that the brain predicts about a category–i.e. the specific features that represent this category in memory. We argued that typical decoders trained to discriminate the bottom-up contrast between two categories (or stimuli) do not access the top-down reactivations of the category-specific features. To address this, we considered classifiers that decode the specific visual features underlying each category–i.e. here, Voltaire and Nuns Gabor features–and then used these to quantify the reactivation of the representations of these specific visual features at Prediction. We showed that top-down reactivations of the category-features (vs. category-contrast) are more precisely localized (i.e. lateralized), more specifically driven by the cues, with per-trial predicted reactivation strength that more strongly biases participant’s perceptual categorization behavior across a wider range of stimulus evidence. Thus, predictions of category-relevant features disentangle the typical category-contrast to improve a mechanistic understanding of prediction processes in the brain, from the top-down cue-reactivation of category-feature representations to the trial-by-trial effect of these reactivations on behavior.

Previous research has shown that a category-contrast can be predicted across the visual hierarchy, from prefrontal cortex to primary auditory and visual sensory areas^17,35^. However, we demonstrated why it is critical to understand the specific category-features that are predicted. Category-contrast classifiers showed bilateral reactivations of the predictions. Instead, we showed that classifiers of the specific category features lateralize the predictions of Voltaire Gabor features in left FG, LG, PCAL and LOC and of Nuns Gabor features in right LOC. Importantly, this LSF/HSF lateralization is consistent with other studies^33,34^ and stimuli and with the Voltaire/Nuns lateralization of our own bottom-up localizer task.

Generalizing from these insights, it is critical to characterize the predicted features because prediction mechanisms will likely align them with the brain’s representations of the input^3,36^, which could change according to the considered region of the occipito-ventral hierarchy. Consider that, in early visual cortex, input representations have known properties of contralateralization, retinotopic mapping^37^ and sensitivity to scale, orientation, and luminance contrast^38,39^; in rFG, input representations are instead more tolerant to changes of scale and orientation^40–43^. Future studies that control stimulus features could therefore parametrically change the scale or the orientation features of the same 3D objects to investigate whether cueing specifically these features (in addition to the object category) reactivates category-features that are scale- and orientation-invariant in rFG, but scale- and orientation-dependent in early visual cortex^44^.

We know that predictions of stimulus information can influence subsequent behavior^7,45^. Advances in neuroimaging further demonstrated that these predictions can modulate the neural responses to inputs by suppressing responses of sensory cortices to predicted stimuli^46–50^ and by modulating premotor cortex activity^51,52^. However, a direct relationship between the strength of the predicted contents in the brain and decision behavior was missing. Previous modelled the single-trial relationship between MVPA representational strength and categorization response time^53^. For the first time, we established here the direct relationship between single-trial prediction strength and perception. Our category-feature classifiers directly quantified the effect of reactivating perception-related category-features on decision behavior bias, whereby stronger feature reactivations led to stronger decision bias across a wider range of feature evidence, all compared with typical category-contrast reactivations. This relationship between reactivation strength and bias was computed for each cue separately, showing that the neural reactivations provide additional information about response bias–i.e. over and above the strong effect of the behavioral cue. Additionally, our results suggest that the prediction bias is stronger with more ambiguous stimuli (i.e., those closer to the PSE). This is consistent with the trade-off effect between prediction and ambiguity of the evidence^8^, whereby observers rely more strongly on predictions when the evidence is ambiguous.

We performed analyses within each participant. To fairly compare the experimental design, we didn’t exclude any participants from the prevalence analysis and reported high probabilities of within-participant replication (7-10 participants, out of 10). However, the behavioral results indicate that Participants 3 and 9 (cf. Supplemental Figure S1) were not engaging in the task correctly–they did not categorize their perception to the presented stimuli, but only responding according to the cue. We further found all the other 8 participants performed the task properly replicated the effect that category-feature reactivations at prediction bias decision behavior.

We traced prediction-for-perception using a realistic and well-known ambiguous stimulus and its two perceptual categorizations. Building from previous research^21,30,54^, we trained participants on using the specific features that enable each perceptual categorization. This approach straightforwardly generalizes to more naturalistic face (e.g. face identity, gender or expression), object (e.g. cars, chairs and buildings) or scene (e.g. indoor rooms, outdoor environments) categorizations. Methodologies already exist to identify the stimulus features that the participant processes to categorize faces^55–57^, objects^58^ and scenes, including when the same input (e.g. a scene image) is categorized in several different ways (e.g. a city, or New York)^51,59^. The key challenge to model predictions is therefore to first identify the specific stimulus features that the participant has represented in memory for each categorization (a cognitive task in the mind of the participant) of a stimulus category (the physical extension of the task, often in the mind of the experimenter). This point brings up the complex question of what the memorized stimulus features are and how we can embed them into a generative model of the stimulus, so we can psychophysically model their dynamic predictions in the brain (see ^56,60–65^ for discussions) as we started developing here.

### Concluding remarks

A mechanistic understanding of prediction-for-perception must quantify where, when, and how brain networks predict the specific category features of the upcoming stimulus to facilitate behavior. With a methodology that can be extended to multiple face, object, and scene categories and across sensory modalities, we showed with increased mechanistic resolution how brain networks predict perceptual contents that directly influence decision behavior.

## Methods

### Participants

Ten participants (18-35 years old, mean=25.1, SD=3.4) took part in the experiment and provided informed consent. All had normal or corrected-to-normal vision and reported no history of any psychological, psychiatric, or neurological condition that might affect visual or auditory perception. The University of Glasgow College of Science and Engineering Ethics Committee approved the experiment (Application Number: 300190094).

### Stimuli

At the Prediction Stage (see Figure 1B), a different pure tone cued Voltaire, Nuns, or had no predictive value (described in Cue-feature training below). The Categorization Stage showed a different hybrid image on each trial. We detail these stimuli below.

#### Prediction Stage: Auditory cues

Pure tones were played for 250 ms, with auditory frequencies of 196Hz (cueing Voltaire), 1760Hz (cueing Nuns) or 880Hz (no prediction).

#### Categorization Stage: Hybrid stimuli

We cropped a grey-level copy of Dali’s *Slave Market with the Disappearing Bust of Voltaire* to retain the ambiguous part of the image that simultaneously shows the bust of Voltaire and the two Nuns across different image scales (256 × 256 pixels, Figure 1A, Original). To extract scale information (i.e. Spatial Frequencies, SF), we decomposed the ambiguous image in 5 bands at 128 (22.4), 64 (11.2), 32 (5.6), 16 (2.8), 8 (1.4) cycles per image (c/deg of visual angle), with brightness set at 0.55.

To quantify the image pixels that underlie selective perceptions of Voltaire and Nuns, we computed the Mutual Information (MI) between the pixel visibility each SF band and perceptual decisions, using data from a previous study^21^. Specifically, we computed MI(Voltaire vs. Don’t Know; pixel visibility) and MI(Nuns vs. Don’t Know; pixel visibility), FWER-corrected over all pixels, *p*<0.05, one-tailed. We found significant pixels at 8 cycles/image for Voltaire and at 32 and 16 cycles/image for Nuns.

To synthesize Hybrid stimuli, we Gabor filtered these significant Voltaire and Nuns pixels to represent them with Gabor coefficients at 6 orientations (0, 30, 60, 90, 120, 150 deg.). Then, we randomly sampled X% of the Voltaire Gabor coefficients and independently Y% of the Nuns Gabor coefficients (2% resolution), while randomly and independently choosing the X and Y percentages according to the distributions shown in in Figure 1A. Finally, we added the X% Voltaire to the Y% Nuns coefficients and shuffled the Gabor coefficients across the background image. We preserved the original contrasts of the Voltaire, Nuns and noisy background Gabors and set their brightness to the same 0.6 value.

### Procedure

#### Auditory localizer

Prior to the main experiment, we ran a MEG localizer session to model the bottom-up processing of each auditory cue. Each trial started with an 250ms pure tone, followed by a 750ms ITI blank. In each 12-trials block, 10 presented the same primary tone; the remaining two tones were catch trials. We instructed participants to press a key whenever they heard a catch tone. Each participant completed 48 such blocks of trials, repeated 16 times for each primary tone, for a total of 576 trials (160 trials per tone).

#### Visual localizer

Prior to the main experiment, we ran another MEG localizer to model the bottom-up processing of 100% Nuns and 100% Voltaire visual features. Each trial started with a 250ms image (with 100% Nuns features, 100% Voltaire features on a mid-grey background) followed by a 1s ITI blank screen. In each 11-trials block, 10 presented the same primary features (e.g. of Voltaire); the remaining image was a catch trial (e.g. of the Nuns). We instructed participants to press a key whenever they saw a catch image. Each participant completed 48 such blocks (i.e., 24 blocks per primary image), for a total of 528 trials (240 trials per primary image).

#### Cue-feature training

Following completion of the localizer tasks, we trained each participant to learn the association between the auditory cues for Voltaire and Nuns and the 100% Voltaire and Nuns features. Each trial started with an auditory cue, followed by a 1s blank screen and an image (100% Nuns or Voltaire) that they categorized as “Voltaire” or “Nuns” (2AFC), followed by feedback (correct vs. incorrect). All participants achieved over 95% accuracy in 75 trials, while implicitly learning the coupling between cues and perceptions. In a second training phase, participants heard an auditory cue and chose the corresponding image (amongst 100% Nuns vs. 100% Voltaire vs. blank image, 3AFC task), followed by feedback (correct vs. incorrect). All participants achieved >90% accuracy in 36 trials.

#### Cueing experiment (Figure 1B)

##### Prediction Stage

Each trial started with a 250ms pure tone (Voltaire, 196Hz; Nuns, 1,760Hz; neutral, 880Hz, each presented on 1/3 of all trials), followed by a 1s blank screen.

##### Categorization Stage

Started with a 500ms fixation followed by a Hybrid image on the centre of the screen that remained until response. Participants responded “Voltaire” vs. “Nuns” vs. “Don’t know” as quickly as they possibly could (3-AFC). A 750ms to 1.25s inter-trial interval (ITI) with jitter followed response.

Each participant completed 45 blocks of 75 such prediction-then-categorization trials, in 4-5 sessions run over 4-5 days, for a total of 3,375 trials per participant.

### MEG Data Acquisition and Pre-processing

We measured each participant’s MEG activity with a 306-channel Elekta Neuromag MEG scanner (MEGIN) at a 1,000Hz sampling rate. We performed the analyses according to recommended guidelines using the MNE-python software^66,67^ and in-house Python/MATLAB code.

We rejected noisy channels with Maxwell filtering and visual inspection, and blocks of trials with a head movement > 0.6cm (tracked by cHPI measurement). For each remaining block, we applied signal-space separation (SSS)^68,69^ to the raw data to reduce environmental noise and compensate for head movement. We band-pass filtered the data between 1-150Hz (Hamming FIR filter), notch-filtered them at 50, 100 and 150Hz and rejected muscle artifacts with automatic detection. We epoched the output data into [-200ms to 2.2s] trial windows around cue onset (visual stimulus onset is at 1.75s), and rejected jump artifacts with automatic detection. We concatenated the epoched data of all blocks per session (i.e., ∼10-16 blocks/day), decomposed the output dataset with ICA, identified and removed the independent components corresponding to artifacts (eye movements, heartbeat—i.e., 2-5 components/participant).

We resampled the output data at 250 Hz, low-pass filtered them at 25Hz (5^th^ Hamming FIR filter) and performed the minimum-norm estimate (MNE) analysis with an empty-room recording. We reconstructed the time series of MEG sources on a 5mm grid of boundary element model (BEM) surface (computed with Freesurfer and MNE software per participant). We applied this reconstruction to each session of trials.

These computations produced for each participant a matrix of single-trial MEG response time series–of dimensions 8,196 MEG sources x 250Hz sampling rate. Supplemental Table S4 reports the number of trials per participant that remained following pre-processing and MNE analysis.

We applied the same pre-processing pipeline to the MEG localizer, using the epoched data [0 to 1000ms] following auditory and visual stimulus onset.

### Analyses

#### Cued-Predictions bias perceptual decisions

To reveal the influence of prediction on decision behavior, we computed the relationship between cues, stimulus feature evidence and perceptual decision probabilities. Pooling all trials across all participants, we quantified the stimulus feature evidence presented on each trial as the difference between its percentage of Voltaire and Nuns Gabor features. We then binned the trials by levels of feature evidence (from -100% to 100%, with 10% steps, Figure 1C, X axis) and calculated “Voltaire,” “Nuns” and “Don’t Know” response probabilities (Figure 1C, Y axis), for each cue type (i.e. Voltaire, Neutral and Nuns panels of Figure 1C). We regressed the feature evidence and the decision probabilities with local linear Gaussian kernels. We computed the Point of Subjective Equality (PSE) as the level of stimulus evidence (i.e. of Voltaire% - Nuns% feature evidence) for which Voltaire and Nuns decisions are equiprobable (Figure 1C).

#### Category-contrast decoding of predictions

##### Localizer cross-validation

To model the bottom-up representations of the auditory cues and of the visual Gabor features, we trained different linear classifiers^70^ (Linear Discriminant Analysis, LDA) using the MEG localizer sensor data. That is, separately for the auditory and visual localizers, we randomly segmented each participant’s MEG trials into 5 folds (without repetitions) and performed a 5-fold cross-validation.

In each validation iteration, we proceeded in 2 steps:

*Step 1: Training*. We trained linear classifiers^70^ (with MNE decoding module^66^) to discriminate the Voltaire vs. Nuns auditory cue and the 100% Voltaire vs. Nuns Gabor features, every 4ms between 0 and 400ms post stimulus, using as training set the MEG sensor data from 4 folds.

*Step 2: Validation*. We tested these trained classifiers every 4ms with the left-out fold. At each time point, we computed classifier decision value as the inner product of the learned linear weights with the held-out fold sensor data.

Following all 5 iterations, we proceeded to Step 3:

*Step 3: Cross-validation performance*. To quantify decoding performance at each time point on a common scale, we concatenated trials across folds and calculated every 4ms the MI^71^ between this classifier decision value and the true stimulus label (i.e. auditory cues for Voltaire vs. Nuns in the auditory classifiers; 100% Voltaire vs. Nuns Gabor features in the visual classifiers). MI quantifies the discrimination information about the stimulus that is available from the classifier weights, without forcing a discrete classification^72^.

We repeated Steps 1 to 3 three times and averaged the resulting three MI matrices (of 100 training time points ×100 testing time points) to quantify the cross-validated decoding performance. To establish statistical significance, we repeated this procedure 1,000 times with shuffled stimulus labels. We applied Threshold-Free Cluster Enhancement^73^ (TFCE, *E*=0.5, *H*=0.5), thresholding for significance with the 95^th^ percentile of the distribution of 1,000 maximum values, each taken across all training × testing time points in the temporal generalization matrix of each shuffle after TFCE (FWER corrected over 100 localizer training time points and 100 localizer testing time points ^74^, *p*<0.05, one-tailed). The resulting matrices of significant MI comprise the time points with significant cross-validation performance. We repeated this cross-validation independently for each participant.

##### Category-contrast reactivation

We used category-contrast classifiers to compute temporal generalization cross-decoding^75^ on the bottom-up processing of the auditory cues contrast and the top-down cued-reactivations of the Voltaire vs. Nuns visual contrast as follows (explained with visual classifiers):

*Step 1: Training*. We trained Voltaire vs. Nuns classifiers on the MEG localizer data at time points of significant cross-validation (see <u>Localizer, cross-validation</u>) between 0 and 400ms post stimulus onset.

*Step 2: Testing*. At the Prediction Stage, every 4 ms between 0 and 600ms post auditory cue, we computed the singular classifier decision value from single-trial MEG sensor response. This produced a 2D (training × testing time) matrix of decision values from the category-contrast classifiers, where each value indicates the reactivation strength of the category contrast at this time point.

*Step 3: Reactivation performance quantification*. For each training x testing time combination, we computed across trials the MI between decision values and ground truth stimulus category (i.e. Voltaire vs. Nuns Gabor features). Permutation testing (1,000 repetitions) established statistical significance (corrected for multiple comparisons with TFCE, FWER corrected over 100 localizer training time×150 Prediction Stage testing time points, one-tailed, *p* < 0.05).

*Step 4: Early vs. Late classifiers*. We split the localizer classifiers into the Early (trained 0-150ms post-stimulus) and Late sets (trained 150-280ms post-stimulus). To measure performance, we chose the Early classifier and the Late classifier with maximum prediction reactivation performance. As above inference was FWER corrected over all classifiers considered.

We repeated Steps 1 to 4 to generate the performance curves of each participant. Figure 3A averages them across participants, separately for Early and Late category-contrast classifiers, for the auditory cues (indicating cue processing) and the visual category contrast (indicating prediction reactivation).

##### Source representation of category-contrast

To localize the MEG source activity underlying auditory and visual category-contrast performance, we must be careful with direct interpretation of weight vectors in sensor space^76^. To address this we used a correlation forward model, where we computed MI between the classifier decision value and source activity, to determine the contribution of each source to the classifier performance^77^. We proceeded as follows:

*Step 1: Time selection*. We selected the two time points of the Prediction Stage when classifier performance peaks–i.e. one in Early prediction (before 140ms post cue); one in Late prediction (after 140ms post-cue).

*Step 2: Source representation reconstruction*. At Early and Late time points, for all 8,196 sources, we computed MI between single-trial category-contrast classifier decision values and single-trial source activity.

We repeated this two-step analysis in each participant to reconstruct their source representations of the auditory and visual category-contrasts at Early and Late Prediction time points. Figure 3B shows their group average; Supplemental Figure S3 shows individual participant results.

#### Category-feature decoding of predictions

##### Category-feature classifier cross-validation

To separately model the dynamic bottom-up representations of Nuns features and Voltaire features, we trained classifiers^70^ (using the MNE decoding module) at the Categorization Stage under neutral cueing. Specifically, every 4ms post-stimulus, we trained binary category-feature Voltaire classifiers on sets of trials with >70% Voltaire (i.e., the top 30% of the trial distribution) vs. <30% Voltaire (i.e., the bottom 30%), and category-feature Nuns classifiers on sets of trials with >70% Nuns vs. <30%. We segmented the participant’s trials into 5 folds based on stratified sampling and performed a 5-fold cross-validation.

In each iteration, and separately for Voltaire and Nuns classifiers, we proceeded as follows:

*Step 1: Training*. We trained linear classifiers to discriminate >70% Voltaire vs.<30% Voltaire, every 4ms between 0 and 400ms post stimulus at the Categorization Stage, under neutral cueing, using MEG sensor data from 4 folds as the training set. We repeated training for the >70% Nuns vs. < 30% Nuns classifiers.

*Step 2: Validation*. We tested the trained classifiers every 4ms on the left-out fold, computing at each time point the decision value as the inner product between classifier weights and held-out fold trials.

Following all 5 iterations, we proceeded to Step 3:

*Step 3: Cross-validation performance*. To quantify decoding performance every 4ms on a common scale, we concatenated trials across folds and calculated MI between this classifier decision value and the true stimulus label (>70% Voltaire/Nuns vs. <30% Voltaire/Nuns).

To quantify cross-validation, we repeated Steps 1 to 3 three times and averaged the resulting three MI matrices (of 100 training ×100 testing time points). We established statistical significance with permutation testing (1,000 repetitions), corrected for multiple comparisons with TFCE–FWER over 100 Categorization Stage training time points × 100 Categorization Stage testing time points, one-tailed, *p* < 0.05. The resulting matrices of significant MI comprised the time points with significant cross-validation performance. We repeated this cross-validation independently for each participant.

##### Category-feature reactivation

We used the category-feature classifiers to separately cross-decode reactivations of Voltaire and Nuns at the Prediction Stage. We traced their source representations as follows:

*Step 1: Training*. We trained the >70% vs. <30% category-feature Voltaire and category-feature Nuns classifiers at time points when they significantly cross-validate (see <u>Category-feature classifier cross-validation</u>), using neutral-cued trials with >70% and <30% Voltaire and Nuns MEG responses.

*Step 2: Testing*. Every 4ms between 0 and 600ms of the Prediction Stage, we tested category-feature classifiers performance on the MEG sensor data. Each trial provides a 2D (training × testing time) matrix of decision values that quantify the reactivation strength of the category-feature prediction.

*Step 3: Reactivation performance quantification*. Separately for Voltaire and Nuns reactivations, and for Voltaire-cued, Nuns-cued and neutral-cued trials, we computed for each training x testing time cell of the matrix the MI between the category-feature classifier values and the ground truth cue labels (i.e. Voltaire, Nuns and neutral). We established statistical significance with permutation testing (1,000 repetitions) and corrected for multiple comparisons with TFCE–FWER over 100 Categorization Stage training time points × 150 Prediction Stage testing time points, one-tailed, *p* < 0.05. We then selected the Voltaire classifier and the Nuns classifier with maximum reactivation performance for the Voltaire Gabor features and separately for the Nuns Gabor features.

*Step 5: Source representations*^76,77^. To visualize the predictions of Voltaire- and Nuns-specific features on MEG sources at their peak reactivation during the Prediction Stage, we computed MI between single-trial category-specific Gabor feature classifier values and MEG source activity. We applied this analysis on all 8,196 sources covering the whole brain.

We repeated Steps 1 to 5 for each participant to produce MEG source representations of Voltaire and Nuns category-feature reactivations (see Figure 4A for their group averages, Supplemental Figure S4 for per-participant results).

#### Specificity of cued-reactivations

To investigate how selectively the cue (for Voltaire and Nuns) reactivates predictions, in we compared per participant the reactivation performance of the category-feature and the category-contrast classifiers as follows:

*Step 1: Classifier selection*. For each participant, across all training (0-400ms post visual stimulus) and testing time points (0-600ms post auditory cue), we selected the three classifiers with maximal reactivation performance: the Voltaire and the Nuns category-feature classifiers (see <u>Category-feature reactivation</u>) and the category-contrast Voltaire vs. Nuns classifier (see <u>Category-contrast reactivation</u>).

*Step 2: Reactivation performance quantification*. We compared these three classifiers on their classification of Prediction Stage MEG sensor data, every 4ms between 0-600ms post-cue (Figure 4B).

#### Reactivation biases behavior

To compare how the trial-by-trial category-feature decoding and category-contrast reactivation at Prediction change response probabilities at Categorization, we examined how their strong vs. weak reactivations change the psychometric relationship between stimulus evidence and response probabilities. For each participant, and separately for Nuns-cued and Voltaire-cued trials, we proceeded as follows:

*Step 1: Time selection*. Every 4ms of the Prediction Stage, we computed the difference of reactivation strength (i.e. classifier decision values) when the participant then categorizes the stimulus as “Voltaire” and as “Nuns” (e.g. on Voltaire-cued trials). We established statistical significance with permutation testing (1,000 repetitions), corrected for multiple comparisons over training × testing time, two-tailed, *p*<0.05. We extracted the single-trial reactivation strength when this significant difference is maximal.

*Step 2: Reactivation split*. We binned the Gabor feature evidence from -100% to 100% (10% step). In each bin, we selected trials with top vs. bottom 30% reactivation strength and compute the probabilities of “Voltaire,” “Nuns,” and “Don’t Know” responses.

We repeated Steps 1 to 2 in each participant and computed the group median of decision probabilities for each bin of feature evidence We then regressed the feature evidence and the group-median decision probabilities (local linear Gaussian kernel), separately for the top and bottom 30% of the trial distribution. Figure 5A and 5B compare these psychometric relationships between category-feature and category-contrast reactivations.

## Acknowledgements

We thank Christoph Daube and Lukas Snoek for comments on earlier versions of this manuscript. This work was funded by the Wellcome Trust (Senior Investigator Award, UK; 107802) and the Multidisciplinary University Research Initiative/Engineering and Physical Sciences Research Council (USA, UK; 172046-01), awarded to Philippe. G. Schyns; and the Wellcome Trust [214120/Z/18/Z], awarded to Robin A.A. Ince. The funders had no role in study design, data collection and analysis, decision to publish or preparation of the manuscript.

