## Supplemental Materials for "Neural representation strength of predicted category features biases decision behavior"

**Supplemental Table S1.** PSE shifts for each participant and type of cue.

| **Participant** | **1** | **2** | **3** | **4** | **5** | **6** | **7** | **8** | **9** | **10** |
|---|---|---|---|---|---|---|---|---|---|---|
| Voltaire cue | 35% | 18% | –↑ | 31% | 28% | –↑ | -13% | 0% | –↑ | –↑ |
| Nuns cue | – ↓ | -33% | – ↓ | – ↓ | – ↓ | – ↓ | -15% | -4% | -46% | 2% |
| Neutral cue | -9% | -17% | -25% | -3% | -25% | -5% | -30% | -7% | 8% | 68% |

*–↑/– ↓ indicates the cases where the curves of Voltaire and Nuns response did not intersect (i.e., PSE out of the stimulus limitation). Specifically, –↑ vs. – ↓ represents that the tendency of PSE shift is towards higher Nuns vs. Voltaire evidence.

**Supplemental Table S2.** PSE shifts for each participant and strong vs. weak cued-reactivations of category-feature (x indicates participants who whom the fit failed)

| **Participant** | **1** | **2** | **3** | **4** | **5** | **6** | **7** | **8** | **9** | **10** |
|---|---|---|---|---|---|---|---|---|---|---|
| Strong Voltaire | **32%** | **30%** | –↑ | **36%** | **36%** | **–↑** | **-1%** | **2%** | –↑ | –↑ |
| Weak Voltaire | 27% | 19% | –↑ | 25% | 21% | 24% | -10% | -5% | –↑ | 37% |
| Strong Nuns | **-30%** | **-36%** | – ↓ | **– ↓** | **– ↓** | – ↓ | **-23%** | **-6%** | – ↓ | **-12%** |
| Weak Nuns | -28% | -26% | – ↓ | 30% | 28% | – ↓ | -10% | 5% | – ↓ | 20% |

*–↑/– ↓ indicates the cases where the curves of Voltaire and Nuns response did not intersect (i.e., PSE out of the stimulus limitation). Specifically, –↑ vs. – ↓ represents that the tendency of PSE shift is towards higher Nuns vs. Voltaire evidence.

***BOLD** indicates statistically significant (FWER-corrected over 100 Categorization training time and 150 Prediction testing time, $p$<0.05, one-tailed) PSE increase in the comparison between strong vs. weak category-feature.

**Supplemental Table S3.** PSE shifts for each participant and strong vs. weak cued-reactivations of category-contrast

| **Participant** | **1** | **2** | **3** | **4** | **5** | **6** | **7** | **8** | **9** | **10** |
|---|---|---|---|---|---|---|---|---|---|---|
| Strong Voltaire | 32% | 22% | –↑ | **46%** | **43%** | 36% | -17% | **2%** | –↑ | 40% |
| Weak Voltaire | 32% | 22% | –↑ | 34% | 20% | –↑ | 0% | -5% | –↑ | x |
| Strong Nuns | **-30%** | -15% | – ↓ | – ↓ | -33% | -40% | -17% | **-10%** | – ↓ | 30% |
| Weak Nuns | -28% | -27% | – ↓ | – ↓ | -34% | -36% | -18% | -6% | – ↓ | – ↓ |

*–↑/– ↓ indicates the cases where the curves of Voltaire and Nuns response did not intersect (i.e., PSE out of the stimulus limitation). Specifically, –↑ vs. – ↓ represents that the tendency of PSE shift is towards higher Nuns vs. Voltaire evidence.

***BOLD** indicates statistically significant (FWER-corrected over 100 Categorization training time and 150 Prediction testing time, $p<0.05$, one-tailed) PSE increase in the comparison between strong vs. weak category-contrast.

**Supplemental Table S4.** Total number of trials per participant following pre-processing and source reconstruction analyses

| Participant | 1 | 2 | 3 | 4 | 5 | 6 | 7 | 8 | 9 | 10 |
|---|---|---|---|---|---|---|---|---|---|---|
| Auditory localizer | 285 | 321 | 303 | 449 | 203 | 419 | 305 | 428 | 421 | 418 |
| Visual localizer | 417 | 455 | 449 | 471 | 352 | 431 | 347 | 378 | 344 | 400 |
| Cueing experiment (Nuns-cued) | 924 | 1065 | 825 | 1045 | 864 | 855 | 1004 | 930 | 872 | 979 |
| Cueing experiment (Voltaire-cued) | 895 | 1074 | 815 | 1035 | 863 | 887 | 1003 | 925 | 917 | 990 |
| Cueing experiment (Neutral-cued) | 927 | 1076 | 800 | 1053 | 842 | 887 | 1016 | 930 | 864 | 977 |

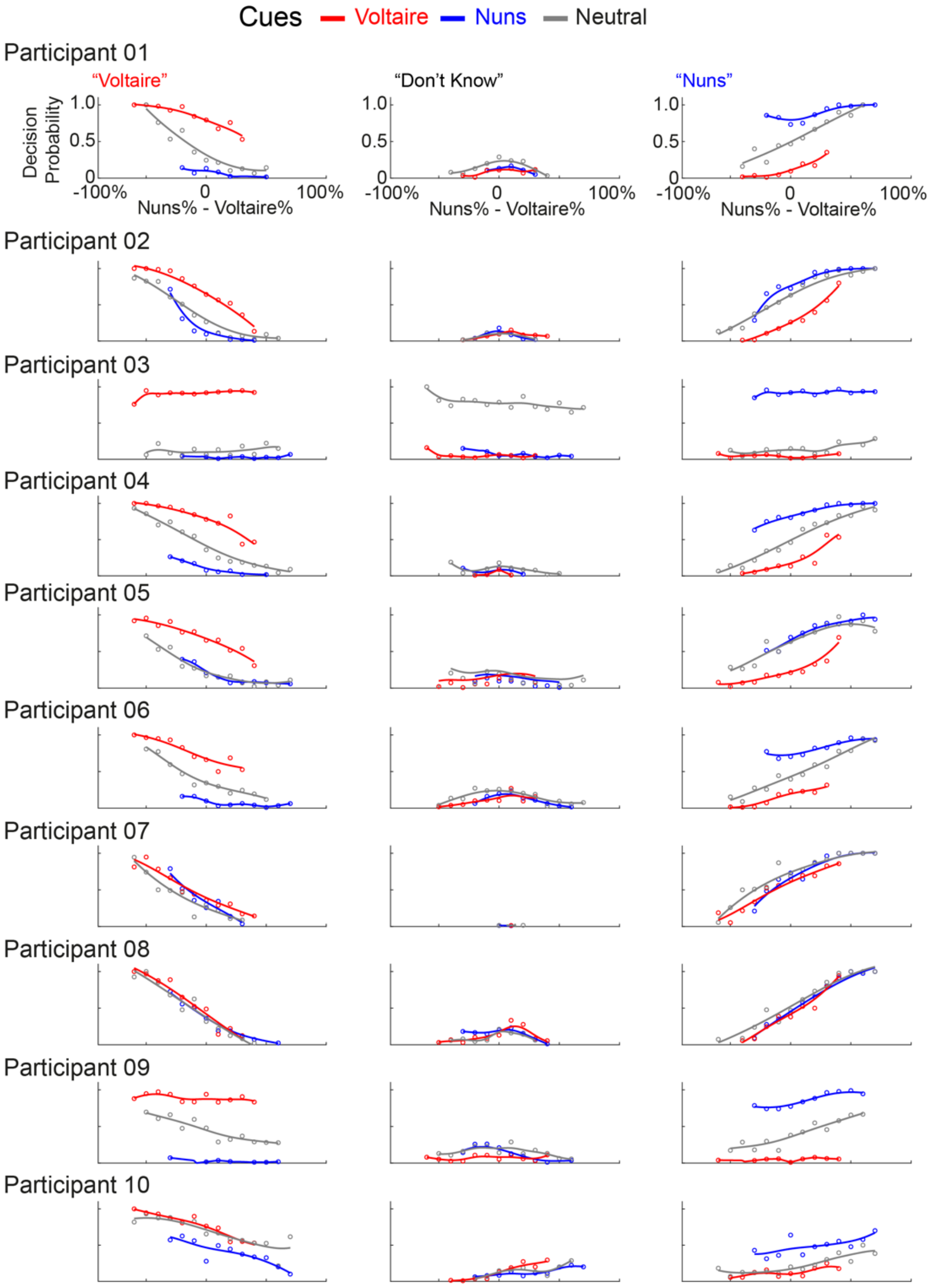


**Supplemental Figure S1. Predictions change individual perceptual decision behavior.** Individual relationships between stimulus feature evidence (X axis) and response probability–i.e. color-coded curves of P("Voltaire"), P("Nuns") and P("Don't Know"), Y axis, for each auditory cue (panel). See the top panel for the labels.

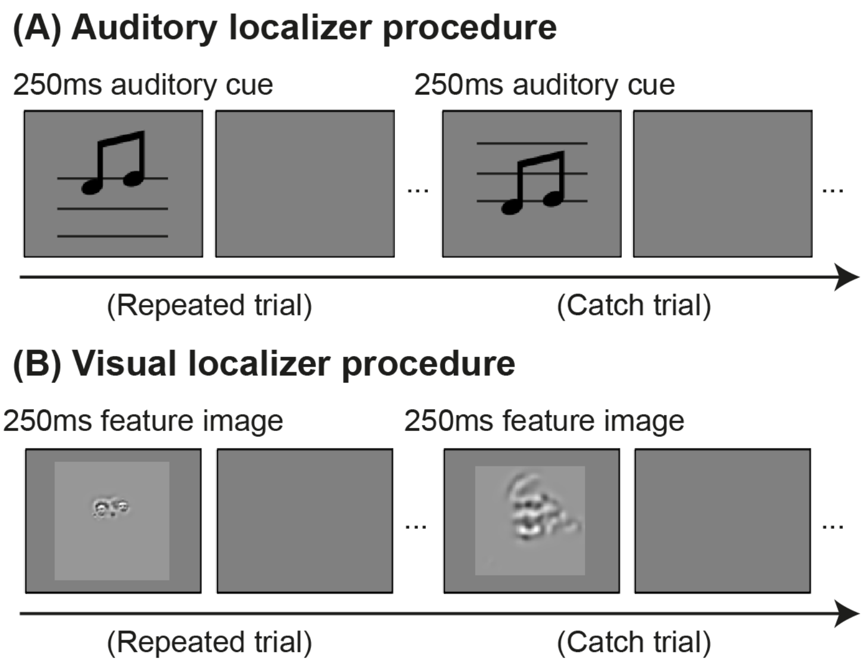


**Supplemental Figure S2. (A) Auditory and (B) visual localizer task procedures.** Prior to the cueing experiment, we ran each participant in an independent auditory MEG localizer (upper panel) and an independent visual MEG localizer (lower panel). In each localizer, each trial started with the 250ms tone or 250ms image followed by a 1000ms ITI blank screen. In each block of 10 trials, participants passively heard 10 repeated tones (i.e., one of 1760Hz, 880Hz, 196Hz) or viewed 10 repeated images (i.e., one of 100% Nuns image, 100% Voltaire image). In the auditory localizer, 2 catch trials displayed two tones with the different frequencies that we instructed participants to detect in each block. In the visual localizer, 1 catch trial presented the image with a different feature.

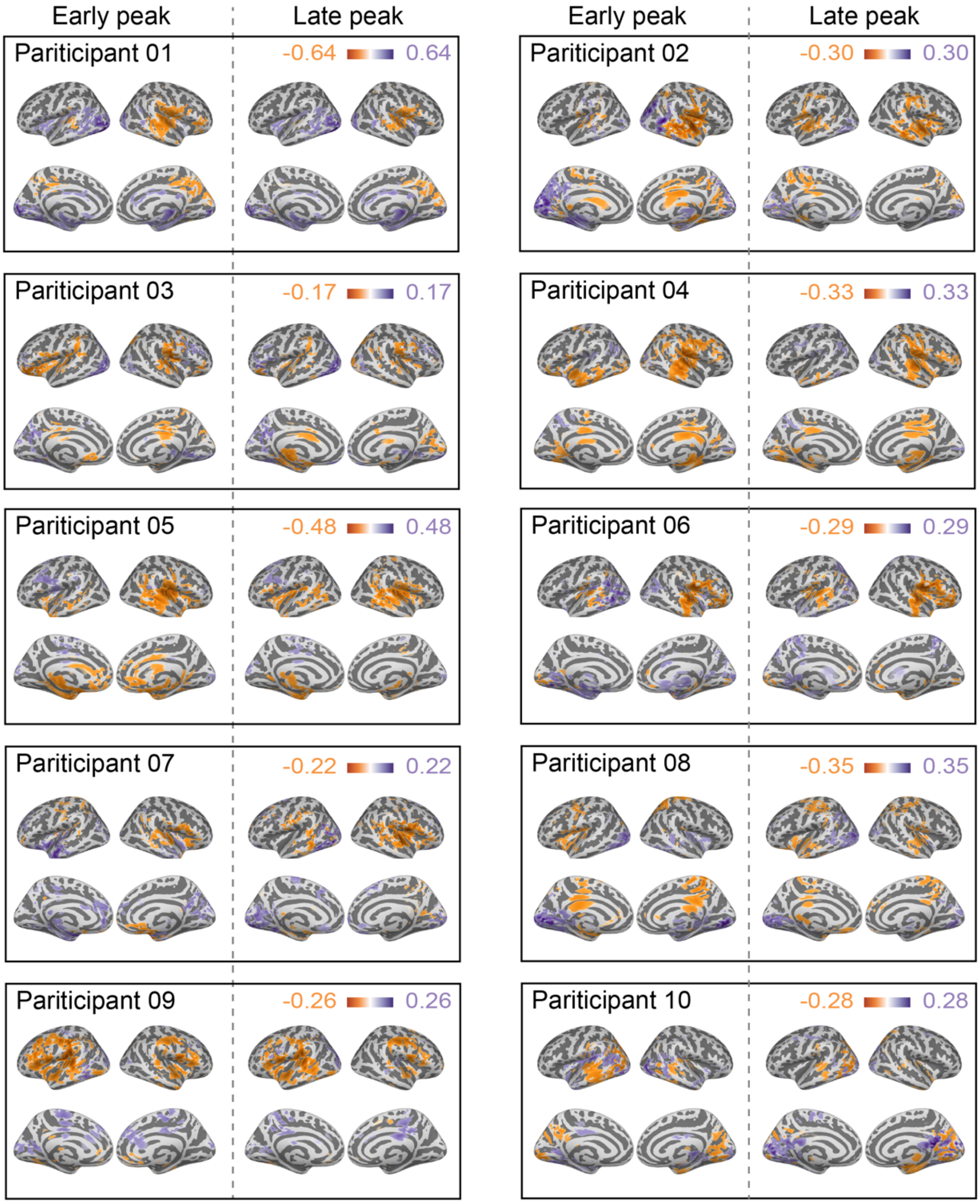


**Supplemental Figure S3. Individual source representations of category-contrast visual reactivation and auditory processing.** We cross-decoded the Prediction Stage using the classifiers trained on the visual localizer (Voltaire vs. Nuns Gabor features) and the auditory localizer (i.e., Voltaire vs. Nuns auditory cues). We localized the MEG sources that contribute to the first (<140ms post-cue) and the second (>140ms post-cue) reactivation performance peaks of the Gabor stimulus features (purple) and auditory cues (orange), computed as MI(decision value of Voltaire vs. Nuns/Gabors or Cues; MEG source activity). The cortical surface plots show the representation difference between visual reactivation and auditory processing for each participant, at the first (left panel) and second (right panel) performance peaks.

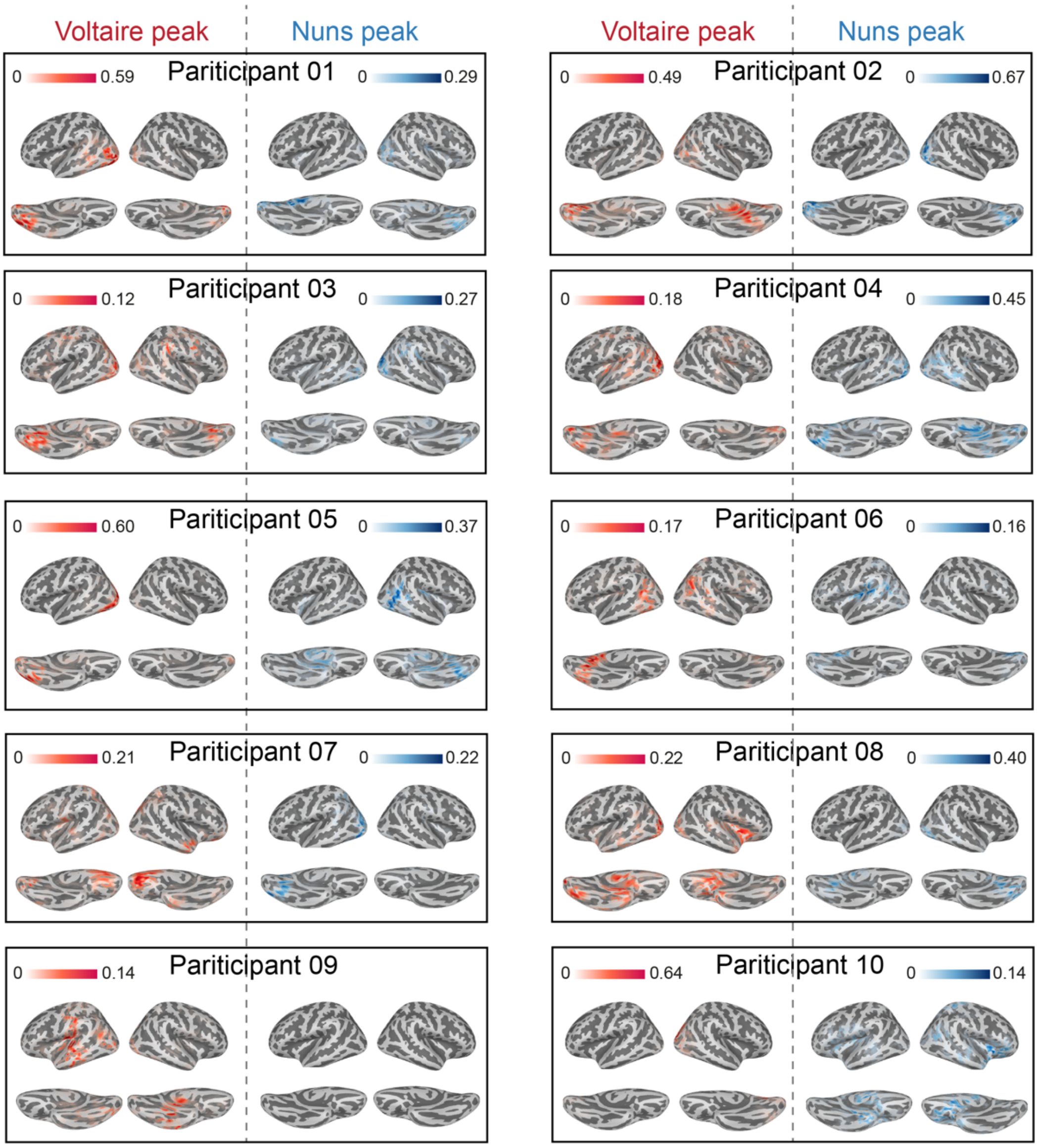


**Supplemental Figure S4. Individual source representations of category-feature reactivation.** We cross-decoded the Prediction Stage with classifiers trained on >70% Voltaire/Nuns vs. <30% Voltaire/Nuns using data from the neutral-cued Categorization Stage. We localized the sources that contribute to the peak reactivation performance, computed as MI(Voltaire/Nuns category-feature reactivation strength; MEG source activity). The cortical surface plots show the representation of category-feature Voltaire (red, left panel) and category-feature Nuns (blue, right panel).
